# RepeatViewer: a platform for explorative analysis of repetitive DNAs

**DOI:** 10.64898/2026.11.23.768287

**Authors:** Jiangzhao Qian, Abdullah El-nagish, Shlok Szatkowski, Sophie Maiwald, Nusrat Sultana, Daniel Costard, Anna Rudzinski, Nicola Schmidt, Tony Heitkam

**Author notes:** These authors contributed equally to this work (Jiangzhao Qian, Abdullah El-nagish).

## Abstract

**Summary:** RepeatViewer is a graphical user interface application designed for repetitive sequence inspection and curation, especially focusing on, but not limited to, transposable elements (TEs). It automatically identifies structural features of TEs, including long terminal repeats (LTRs), terminal inverted repeats (TIRs), and target side duplications (TSDs). All detected and user-defined annotations can be interactively inspected, edited, and exported. RepeatViewer also integrates a TE curation workflow intended to facilitate their manual refinement. In addition, it provides interactive visualization of TE sequences and genome-wide TE annotations, such as dot plots as well as TE distribution and divergence analyses. RepeatViewer is modularly built and can be easily customized by the user.

**Availability and Implementation:** RepeatViewer is implemented in Python (version 3.10 or later) and is freely available at https://github.com/qjiangzhao/RepeatViewer. It runs on Linux, macOS, and Windows and is distributed as a prebuilt package.

## 1 Introduction

Repetitive DNA sequences occur across most taxa and influence genome evolution, genome size, gene expression, and the generation of genetic variation (Bourque et al., 2018; Wells and Feschotte, 2020). Yet, repetitive DNA identification and annotation is complex and error-prone. This applies to both, annotation of transposable elements (TEs) as well as of the tandemly repeated genome fraction. Especially, the precise annotation of TEs is hampered by high TE sequence diversity and the existence of fragmented and nested TE copies. Several softwares, including RepeatModeler2, EDTA, DANTE, REPET4.0 and others, generate TE consensus libraries automatically (Flynn et al., 2020; Ou et al., 2019; Novák et al. 2024, Bardet et al. 2025). However, the output TE consensus sequences are often fragmented, nested, redundant, and with inaccurate boundaries. Therefore, manual curation of the TEs is still required (Storer et al., 2021; Goubert et al., 2022). The manual curation of TEs commonly involves several independent command-line tools and operations, consumes a lot of time, and requires a high level of expertise in the field of TEs. As the human contribution in these analyses is still rather large, reproducibility may suffer. Tools such as TEtrimmer automate many of these procedures and provide a graphical user interface for manual review of the outputs (Qian et al., 2025). However, a general user-friendly graphical platform for the inspection, annotation, and visualization of repetitive DNA sequences and genome-wide annotations is still lacking. Here, we introduce RepeatViewer, a graphical user interface application that integrates these functions to support the repetitive DNA analysis for users of all experience levels.

## 2 Implementation and features

### 2.1 Overview of RepeatViewer

RepeatViewer is developed with Python3 using the PySide6 package (The Qt Company, 2026). It is a cross-platform tool, which can be installed and executed on Linux, macOS, and Windows systems. Its core functions are organized into five modules, including file navigation and preview, sequence utilities, visualization, analysis, and extensibility; the detailed functions and tools under each module are listed in Fig. 1A. The application window contains three panels: (i) the directory panel for navigation, (ii) the file panel for file selection, and (iii) the preview panel enabling the display of sequences and analysis results (Fig. 1B). When displaying a FASTA sequence, three tabs are provided in the preview panel; those are “Text”, “Annotation”, and “Dotplot(self)”. The Annotation tab displays all annotated sequence features.

**Fig. 1.**
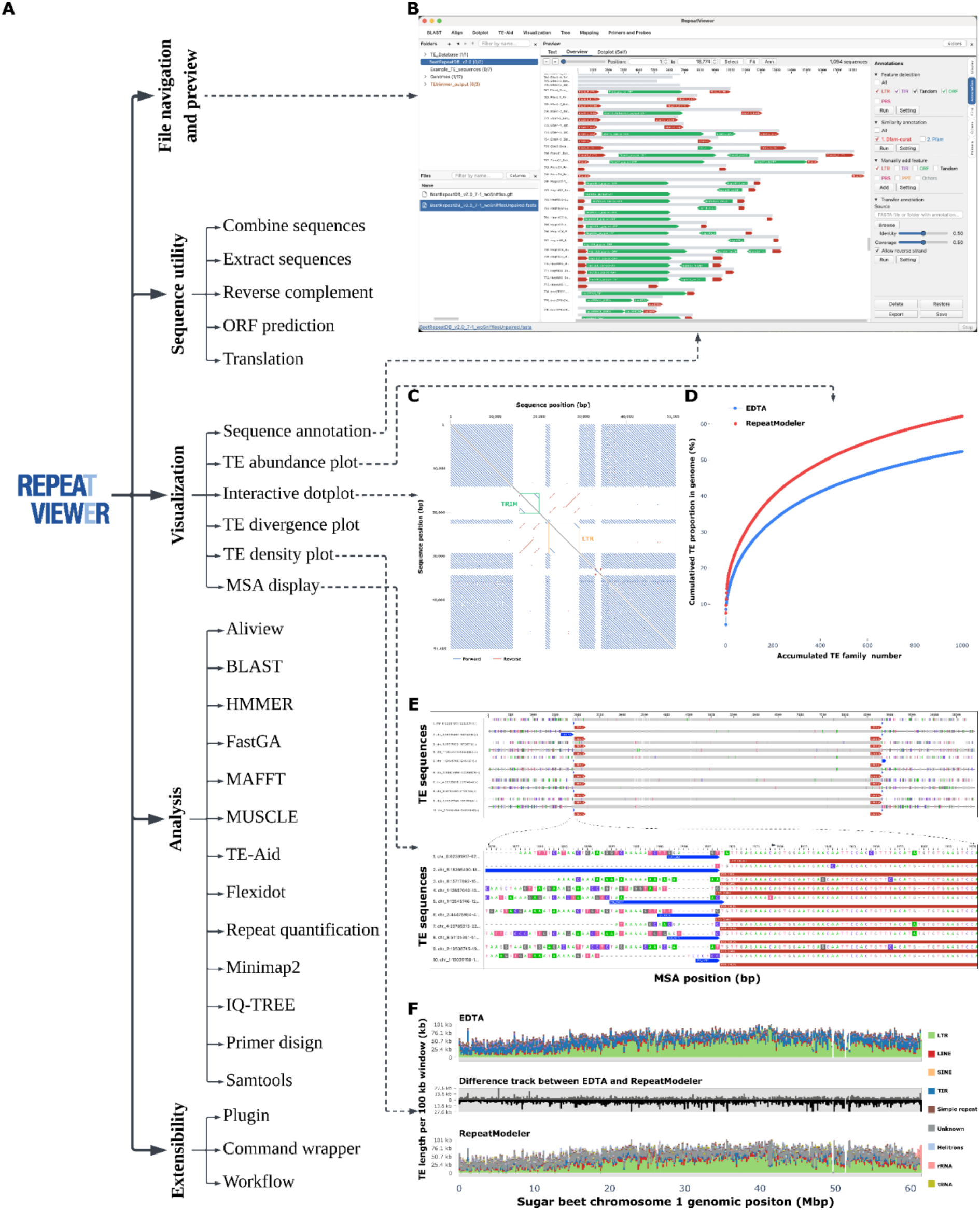
Overview of RepeatViewer interface and functions and example TE visualizations from *Beta vulgaris*. (A) Functional overview of RepeatViewer, grouping file navigation and preview, sequence utilities, visualization, analysis, and extensibility. (B) The graphical user interface of RepeatViewer. The upper-left directory panel shows the working project tree. The lower-left files panel lists contents of the selected directory with sequence metadata. The preview panel on the right opens the selected FASTA file in the interactive sequence viewer. A collection of TE sequences from sugar beet is shown as example (Schmidt *et al*., 2024); predicted LTRs and ORFs are shown as dark red and green arrows respectively. (C) Interactive self-comparison dotplot for the centromeric satellite DNA sequence beetSat01-pBV (Schmidt *et al*., 2024) from *Beta vulgaris* in RepeatViewer. The satellite DNA array is interrupted by full-length copies of a non-autonomous TRIM element and its autonomous partner, the Ty3/*gypsy* LTR retrotransposon Beetle7 (the latter being in reverse orientation). Both axes indicate the position along the sequence in bp. Blue lines represent same strand nucleotide repetitions and red lines are the reverse complement matched nucleotide repetitions. Here, a word size of 14 was chosen. Users can easily zoom and check the sequences for each line. (D) TE abundance plot for RepeatMasker outputs based on EDTA (blue) and RepeatModeler (red) using the *Beta vulgaris* genome. Each dot represents one TE family. Families are sorted by occupied genome proportion from high to low along the x-axis (top 1,000 families shown). The y-axis is the accumulated TE proportion of the genome. (E) RepeatViewer displays TE-derived MSAs with LTR and TSD annotations. The upper panel shows the complete alignment, while the lower panel provides a zoomed-in view of the 5′ LTR region. The x-axis represents MSA position (bp), and the y-axis represents individual TE sequences. (F) TE density plot for *Beta vulgaris* chromosome one. The upper and middle tracks show the total TE amount per 100 kb window for RepeatMasker annotations generated based on the EDTA and RepeatModeler libraries, respectively. TE types are colored differently. The middle track shows the TE differences between the EDTA and RepeatModeler-based TE annotation. Values above zero represent annotated TEs unique to EDTA, and values below zero mark annotated TEs unique to RepeatModeler. The x-axis indicates the genomic position in Mbp.

RepeatViewer can process DNA and protein FASTA files, RepeatMasker output tables (Smit et al., 2013–2015), GFF/GTF/BED/VCF annotations, PAF, BAM alignments, Bismark methylation coverage tracks (Krueger and Andrews, 2011), and user-defined DNA, protein, and HMM databases. Users can import folders by dragging them to the directory panel. All files in the folder can be listed in the file panel by clicking the corresponding folder name. Most of the analysis results are shown in the preview panel (Fig. 1B).

### 2.2 Interactive annotation of TE structure and proteins

For any TE, the detection of representative structural features is needed for TE classification and boundary definition. For example, intact long terminal repeat (LTR) retrotransposons contain two flanking LTRs and many DNA transposons have terminal inverted repeats (TIRs) (Wicker et al., 2007). RepeatViewer quickly identifies LTRs, TIRs, and target site duplication (TSD) for any given TE sequence and TE-derived multiple sequence alignment (MSA). Users can modify feature detection parameters, e.g. minimum and maximum length of the feature, the distance between paired terminal repeats, and detection similarity thresholds. All predicted features are recorded to a GFF file and displayed in the sequence annotation tab colored individually (Fig. 1 B).

Autonomous TEs always encode characteristic proteins for their transposition (Wicker et al., 2007), which are necessary for further classification of different TEs. Using RepeatViewer, open reading frames (ORFs) can be predicted with the python package “orfipy” (Singh and Wurtele, 2021) and indexing is possible with multiple user-defined databases like Dfam (Storer et al., 2021), Repbase (Bao et al., 2015), REXdb (Neumann et al., 2019), Pfam (Mistry et al., 2021), and RepetDB (Amselem et al., 2019) with a single button operation. All hits are displayed together with the structural features in the annotation tab (Fig. 1B) and can be exported in a GFF format. Users can further add, modify, recolor, or delete any annotation features.

### 2.3 TE consensus sequence curation

Manual curation of TEs usually starts with a BLASTn search (Camacho *et al*., 2009) of a consensus or reference sequence against a genome of choice, followed by the extraction and extension of genomic copies, multiple sequence alignment, and TE boundary definition (Goubert et al., 2022; Baril et al., 2024; Orozco-Arias et al., 2024; Qian et al. 2025). RepeatViewer integrates the tools required for this procedure as a workflow. Users can execute the entire procedure by a single button operation, which can decrease the TE curation workload considerably. Besides, users can design their own workflows easily to customize their analysis. Besides, RepeatViewer integrates MSA display and modification functions, so that users can quickly edit TE-derived MSAs for the final TE consensus sequence generation (Fig. 1E).

### 2.4 Visualization of TE sequences and genome-wide TE annotations

RepeatViewer provides several visualization functions for checking individual TE sequences and genome-wide TE annotations. TE-Aid-style plots (Goubert *et al*., 2022) summarize BLASTn hit length and coverage for a selected TE consensus sequence. FlexiDot (Seibt *et al*., 2018) generates self, pairwise, and multi-sequence dot plots. In addition, RepeatView provides an interactive dotplot display (Fig. 1C), enabling users to navigate from a selected plot region to the corresponding sequence position in the annotation tab. At the genome level, RepeatViewer generates TE abundance plots (Fig. 1D) to identify the predominant TE families/lineages, and TE divergence plots to show sequence divergence of genomic TE copies from the respective consensus. Besides, visualization of genomic distribution and TE density within selected genomic regions is enabled by TE density plots (Fig. 1F), which also allows comparison of two annotation results for the same genome. These functions facilitate the visual inspection of TE structure, TE multiple sequence alignments, genomic distribution, and differences between alternative annotations.

### 2.5 Tandem repeat and satellite DNA analysis

Besides interspersed repeats, tandem repeats, from short microsatellites to satellite DNA arrays spanning megabases, make up a substantial fraction of most plant and animal genomes, yet tools for their manual inspection are even scarcer than for TEs.

RepeatViewer analyzes tandem repeats by detecting short, generic and approximate tandem repeats with “pytrf” (Du et al., 2025). Those are written to GFF, allowing inspection and editing of monomer boundaries by hand. Self dotplots generated with FlexiDot resolve the head-to-tail pattern characteristic of tandem arrays, from which monomer length and higher-order organisation can be read directly, and any selected repetition can be traced back to its position in the annotation tab. The curation workflow described in 2.3 applies unchanged to a satellite DNA monomer consensus: a BLASTn search against the genome, extraction and extension of genomic copies, and multiple sequence alignment refine a monomer exactly as they refine a TE. Finally, the Repeat quantification function searches a library of monomer or satellite DNA consensus sequences against a target assembly and reports copy number, number of arrays, occupied base pairs and proportion of the target assembly, and mean identity per monomer family, together with a GFF track of array positions, which can then be displayed alongside TE annotations and DNA methylation coverage in the genome-wide density view.

## 3 Discussion

Accurate repetitive DNA annotations still need manual inspection. Tools such as MCHelper (Orozco-Arias et al., 2024) and TEtrimmer (Qian et al., 2025) can automate the majority of the curation process, but expert checking remains necessary. RepeatViewer facilitates this process with a graphical user interface designed for repetitive sequence analysis, including workflow functions to integrate multiple steps into one button. In practice, for TE analysis, we recommend generating TE libraries with TEtrimmer and then using RepeatViewer to inspect and refine the output. This combination keeps the speed of automation while retaining expert control over TE quality.

Unlike other commercial graphical tools, RepeatViewer is fully free and open source, making repetitive DNA curation more accessible to laboratories without expensive software licenses. Compared with multi-purpose free tools such as UGENE (Okonechnikov et al., 2012), it focuses on TE-oriented tasks, from structural feature annotation and consensus curation to visualization and genome-wide comparison. To support further customization, RepeatViewer provides extensible functions including plugin, command wrapper, and workflow. We hope that RepeatViewer can serve as a shared framework in which the TE community (e.g. those within TE Hub; TE Hub Consortium et al., 2021) contributes and exchanges desired functions, making new tools immediately available to all users.

## Abbreviations

BLAST: Basic Local Alignment Search Tool
CDD: Conserved Domain Database
GFF: General Feature Format
GUI: Graphical user interface
LTR: Long terminal repeat
MSA: Multiple sequence alignment
ORF: Open reading frame
TE: Transposable element
TIR: Terminal inverted repeat
TSD: Target Site Duplication

## Acknowledgements

We thank members of the Chair of Molecular Botany at RWTH Aachen for discussions and testing during the development of RepeatViewer.

## Author contributions

J.Q. and T.H. conceived the study. T.H. provides the funding. J.Q. and A.E. developed RepeatViewer. A.E. developed the RepeatViewer website. S.M. developed the PBS identification module. A.E. and N.Su. drafted the manual. S.S., D.C., A.R., S.M. and N.Sc. contributed to software testing and debugging. All authors wrote, reviewed and approved the manuscript.

## Data availability

The source code and documentation of RepeatViewer are freely available at https://github.com/qjiangzhao/RepeatViewer. A versioned software archive was deposited in Zenodo (DOI: 10.5281/zenodo.24429519). We used coding agents, i.e. Claude code and Cursor, to facilitate the development of RepeatViewer.

